# DNA methylation links prenatal smoking exposure to later life health outcomes in offspring

**DOI:** 10.1101/428896

**Authors:** Petri Wiklund, Ville Karhunen, Rebecca C Richmond, Alina Rodriguez, Maneka De Silva, Matthias Wielscher, Faisal I Rezwan, Tom G Richardson, Juha Veijola, Karl Heinz-Herzig, John W Holloway, Caroline L Relton, Sylvain Sebert, Marjo-Riitta Järvelin

**Affiliations:** Center for Life Course Health Research, University of Oulu, Oulu, Finland; Department of Epidemiology and Biostatistics, Imperial College London, London, UK; Department of Health Sciences, University of Jyvaskyla, Jyvaskyla, Finland; MRC Integrative Epidemiology Unit, University of Bristol, UK; College of Social Sciences, University of Lincoln, UK; Human Development and Health, Faculty of Medicine, University of Southampton, Southampton, UK; Medical Research Center Oulu, Finland; Oulu University Hospital, Oulu, Finland; Research Unit of Clinical Neuroscience, University of Oulu, Oulu, Finland; Institute of Biomedicine and Biocenter of Oulu, Finland; Department of Gastroenterology and Metabolism, Poznan University of Medical Sciences, Poznan, Poland; Clinical and Experimental Sciences, Faculty of Medicine, University of Southampton, Southampton, UK; Department for Genomics of Common Diseases, School of Medicine, Imperial College London, London, UK; MRC-PHE Centre for Environment and Health, Imperial College London, London W2 1PG, UK; Department of Life Sciences, College of Health and Life Sciences, Brunel University London, United Kingdom

**Author notes:** These authors contributed equally to this work (Petri Wiklund, Ville Karhunen). Corresponding author: Marjo-Riitta Järvelin.

## Abstract

Maternal smoking during pregnancy is associated with adverse offspring health outcomes across their life course. We hypothesize that DNA methylation is a potential mediator of this relationship. To test this, we examined the association of prenatal maternal smoking with DNA methylation in 2,821 individuals (age 16 to 48 years) from five prospective birth cohort studies and perform Mendelian randomization and mediation analyses to assess, whether methylation markers have causal effects on disease outcomes in the offspring. We identify 69 differentially methylated CpGs in 36 genomic regions (P < 1×10^−7^), and show that DNA methylation may represent a biological mechanism through which maternal smoking is associated with increased risk of psychiatric morbidity in the exposed offspring.

## Introduction

Maternal smoking during pregnancy is associated with increased risk for pre-term birth, fetal growth restriction and low birth weight^1–3^, as well as neurodevelopmental impairments and respiratory and cardiovascular diseases later in life^4–8^. Despite these well-known risks, many women who commence pregnancy as smokers continue to smoke throughout gestation. According to a recent meta-analysis, the global prevalence of maternal smoking during pregnancy varies widely from a few percentages up to nearly 40% in Ireland^9^. Thus, cigarette smoking continues to be one of the most important modifiable risk factors for the health of mothers and their children.

Cigarette smoke is a potent environmental modifier of DNA methylation^10^. In support of this, an epigenome-wide meta-analysis of 13 birth cohort studies identified over 6,000 differentially methylated CpGs in cord blood of newborns exposed to prenatal smoking^11^. Several smaller studies have suggested that some of these methylation changes may persist across childhood and adolescence^12,13^into adulthood^14,15^. However, questions remain concerning whether such DNA methylation changes endure across the life course and whether they play a mediating role in linking prenatal smoke exposure to later life health outcomes.

Here, we combine data from five prospective birth cohort studies to investigate associations between prenatal smoking exposure and DNA methylation in 2,821 adolescents and adults. We first examine the associations of prenatal smoking exposure with DNA methylation in each cohort and then meta-analyze the results across all studies. We focus on the >6,000 CpG sites previously identified in cord blood of newborns exposed to prenatal smoking^11^. We further (i) assess the impact of current smoking by the participant on DNA methylation; (ii) explore the dose-dependent effects of prenatal smoking exposure on methylation at key CpG sites; (iii) examine the potential intrauterine effect of smoking exposure on offspring DNA methylation by using paternal smoking as a negative control; (iv) assess the persistence of DNA methylation changes by investigating longitudinal associations from 30 to 48 years of age; and (v) conduct Mendelian randomization (MR) and mediation analyses to examine the potential causal effects of DNA methylation changes on disease outcomes in the offspring (Figure 1). Our results show that prenatal smoking has persistent effects on the offspring epigenome and provide evidence for a causal role of DNA methylation in adverse health effects that may arise from exposure to tobacco smoke in utero.

**Figure 1.**
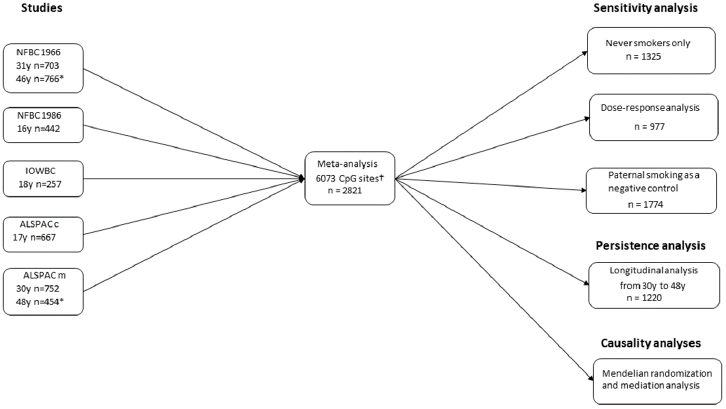
Study design and analytical flow of the study. NFBC = Northern Finland Birth Cohort, ALSPAC = Avon Longitudinal Study of Parents and Children (m=mothers) and (c=children), IWBC = Isle of Wight Birth Cohort, EWAS = epigenome-wide association study. †CpG sites identified previously in cord blood of newborns exposed to maternal smoking in utero (20). *Methylation data for persistence analysis.

## Results

### Cohort specific characteristics of the study participants

We analyzed the association of prenatal smoking exposure with DNA methylation in altogether 1,366 adolescents (age 16y to 18y) and 1,455 adults (age 30y to 31y). Of these, 1,145 were from two independent Northern Finland Birth cohorts (NFBC 1966 and NFBC 1986), 257 were from the Isle of Wight Birth Cohort (IOWBC) and 1,419 from two Avon Longitudinal Study of Parents and Children cohorts (ALSPAC mothers and ALSPAC children). **Supplementary Tables 1-3** show the characteristics of each study cohort. Overall, 18.4% of the NFBC 1966 and 13.2% of the NFBC 1986 were prenatally exposed to maternal smoking. The corresponding figures were 11.8% for ALSPAC children, 28.7% for ALSPAC mothers and 16.3% for IOWBC.

### DNA methylation meta-analysis

We found evidence for 69 differentially methylated CpGs in 36 genomic regions (Table 1). All of these CpG sites showed directionally concordant effects with previously reported associations in newborns^11^ e.g., hypermethylation of cg04180046 in *MYOG1* and cg05549655 in *CYP1A1* and hypomethylation of cg05575921 in *AHRR* and cg14179389 in *GFI1* in the exposed offspring compared with their unexposed counterparts.

**Table 1.** Association of exposure to maternal smoking during pregnancy and offspring peripheral blood DNA Methylation

| CpG | Chr | Position | Gene | $\beta$ (SE) | P |
| --- | --- | --- | --- | --- | --- |
| cg04180046 | 7 | 45002736 | *MYO1G* | 0.045 (0.003) | 2.60E-54 |
| cg12803068 | 7 | 45002919 | *MYO1G* | 0.077 (0.005) | 5.70E-45 |
| cg22132788 | 7 | 45002486 | *MYO1G* | 0.045 (0.003) | 6.80E-38 |
| cg19089201 | 7 | 45002287 | *MYO1G* | 0.035 (0.003) | 1.20E-31 |
| cg05549655 | 15 | 75019143 | *CYP1A1* | 0.010 (0.001) | 1.00E-28 |
| cg25949550 | 7 | 145814306 | *CNTNAP2* | -0.007 (0.001) | 5.40E-27 |
| cg18493761 | 11 | 125386885 |  | 0.037 (0.004) | 7.30E-21 |
| cg15507334 | 10 | 14372913 | *FRMD4A* | 0.024 (0.003) | 1.80E-20 |
| cg14179389 | 1 | 92947961 | *GFI1* | -0.028 (0.003) | 5.00E-20 |
| cg00253658 | 16 | 54210496 |  | 0.037 (0.004) | 3.40E-19 |
| cg17924476 | 5 | 323794 | *AHRR* | 0.024 (0.003) | 4.40E-19 |
| cg13570656 | 15 | 75019196 | *CYP1A1* | 0.036 (0.004) | 7.40E-19 |
| cg11813497 | 10 | 14372879 | *FRMD4A* | 0.026 (0.003) | 9.30E-19 |
| cg22549041 | 15 | 75019251 | *CYP1A1* | 0.041 (0.005) | 2.60E-18 |
| cg05575921 | 5 | 373378 | *AHRR* | -0.019 (0.002) | 3.50E-18 |
| cg12101586 | 15 | 75019203 | *CYP1A1* | 0.032 (0.004) | 1.20E-17 |
| cg18092474 | 15 | 75019302 | *CYP1A1* | 0.036 (0.004) | 1.50E-17 |
| cg11924019 | 15 | 75019283 | *CYP1A1* | 0.013 (0.002) | 2.90E-16 |
| cg25464840 | 10 | 14372910 | *FRMD4A* | 0.019 (0.002) | 9.30E-16 |
| cg00213123 | 15 | 75019070 | *CYP1A1* | 0.012 (0.002) | 7.80E-15 |
| cg11207515 | 7 | 146904205 | *CNTNAP2* | -0.023 (0.003) | 1.00E-13 |
| cg14157435 | 2 | 206628692 | *NRP2* | -0.046 (0.006) | 1.10E-13 |
| cg05348875 | 2 | 206628625 | *NRP2* | -0.040 (0.006) | 7.60E-13 |
| cg01952185 | 5 | 134813213 |  | 0.019 (0.003) | 3.30E-12 |
| cg22308949 | 2 | 206628553 | *NRP2* | -0.022 (0.003) | 4.40E-12 |
| cg05204104 | 2 | 235403141 | *ARL4C* | 0.017 (0.003) | 4.80E-12 |
| cg11641006 | 2 | 235213874 |  | 0.017 (0.003) | 7.90E-12 |
| cg09935388 | 1 | 92947588 | *GFI1* | -0.029 (0.004) | 1.10E-11 |
| cg05697249 | 11 | 111789693 | *C11orf52* | 0.015 (0.002) | 1.30E-11 |
| cg21161138 | 5 | 399360 | *AHRR* | -0.013 (0.002) | 1.50E-11 |
| cg26681628 | 16 | 54210550 |  | 0.018 (0.003) | 1.80E-11 |
| cg11025974 | 2 | 152830521 | *CACNB4* | 0.016 (0.002) | 9.10E-11 |
| cg15016771 | 2 | 235403218 | *ARL4C* | 0.008 (0.001) | 1.00E-10 |
| cg11429111 | 5 | 134813329 |  | 0.014 (0.002) | 1.30E-10 |
| cg25189904 | 1 | 68299493 | *GNG12* | -0.020 (0.003) | 1.30E-10 |
| cg06758350 | 21 | 36259460 | *RUNX1* | 0.030 (0.005) | 1.70E-10 |
| cg17852385 | 15 | 75019188 | *CYP1A1* | 0.010 (0.002) | 2.60E-10 |
| cg01664727 | 21 | 36258423 | *RUNX1* | 0.027 (0.004) | 2.60E-10 |
| cg20344448 | 10 | 14372431 | *FRMD4A* | 0.014 (0.002) | 2.80E-10 |
| cg00794911 | 6 | 166260532 |  | -0.012 (0.002) | 5.20E-10 |
| cg22698744 | 21 | 36263808 | *RUNX1* | 0.023 (0.004) | 7.30E-10 |
| cg03142697 | 21 | 36258497 | *RUNX1* | 0.016 (0.003) | 8.10E-10 |
| cg14563637 | 9 | 98931801 |  | 0.016 (0.003) | 1.20E-09 |
| cg15091747 | 21 | 36262896 | *RUNX1* | 0.014 (0.002) | 1.20E-09 |
| cg12984635 | 19 | 44032076 | *ETHE1* | 0.015 (0.002) | 2.20E-09 |
| cg01825213 | 9 | 98979965 |  | 0.019 (0.003) | 2.50E-09 |
| cg12477880 | 21 | 36259241 | *RUNX1* | 0.038 (0.006) | 2.80E-09 |
| cg17199018 | 8 | 28206278 | *ZNF395* | -0.017 (0.003) | 3.90E-09 |
| cg00174179 | 3 | 49450293 | *RHOA;TCTA* | -0.006 (0.001) | 7.00E-09 |
| cg15578140 | 7 | 147718109 | *MIR548F3;CNTNAP2* | 0.011 (0.002) | 7.50E-09 |
| cg14540913 | 9 | 132458514 | *PRRX2* | 0.013 (0.002) | 7.60E-09 |
| cg18132363 | 6 | 166260572 |  | -0.019 (0.003) | 7.60E-09 |
| cg21253335 | 5 | 87835928 |  | 0.017 (0.003) | 1.30E-08 |
| cg25879142 | 7 | 4671391 |  | 0.018 (0.003) | 1.80E-08 |
| cg11845417 | 11 | 111789613 | *C11orf52* | 0.011 (0.002) | 2.40E-08 |
| cg20117519 | 7 | 8429907 |  | 0.022 (0.004) | 2.40E-08 |
| cg05783384 | 2 | 218843735 |  | 0.021 (0.004) | 2.60E-08 |
| cg13822849 | 9 | 137999757 | *OLFM1* | 0.007 (0.001) | 2.90E-08 |
| cg16449012 | 4 | 17781880 | *FAM184B* | 0.014 (0.002) | 3.10E-08 |
| cg05634495 | 6 | 122364658 |  | 0.016 (0.003) | 3.10E-08 |
| cg08644678 | 4 | 17711202 | *FAM184B* | 0.009 (0.002) | 3.10E-08 |
| cg13834112 | 15 | 90361639 |  | 0.013 (0.002) | 3.60E-08 |
| cg06635952 | 2 | 70025869 | *ANXA4* | 0.011 (0.002) | 5.50E-08 |
| cg14485097 | 7 | 4671479 |  | 0.016 (0.003) | 7.40E-08 |
| cg04598670 | 7 | 68697651 |  | -0.019 (0.003) | 8.30E-08 |
| cg03252786 | 11 | 125106056 | *PKNOX2* | 0.006 (0.001) | 8.70E-08 |
| cg04749740 | 2 | 65935124 |  | 0.015 (0.003) | 9.20E-08 |
| cg15325070 | 1 | 2792704 |  | 0.014 (0.003) | 9.20E-08 |
| cg04358214 | 16 | 67143304 | *C16orf70* | 0.022 (0.004) | 9.60E-08 |

The analyses were conducted separately in each participating cohort adjusted for offspring age, sex, body mass index, smoking status and genetic ancestry, and combined using inverse-variance weighted fixed effects meta-analysis. Chr = chromosome; $\beta$ = effect size estimate, SE = standard error.

### Sensitivity and downstream analyses

To examine whether offsprings’ own smoking had influenced the results, we repeated the main analysis including only those individuals who had never smoked regularly in their life. The results were similar, in both direction and magnitude, across all 36 genomic regions as in the full meta-analysis (Figure 2), indicating that the association between maternal smoking and DNA methylation was not mediated through offsprings’ own smoking behavior.

**Figure 2.**
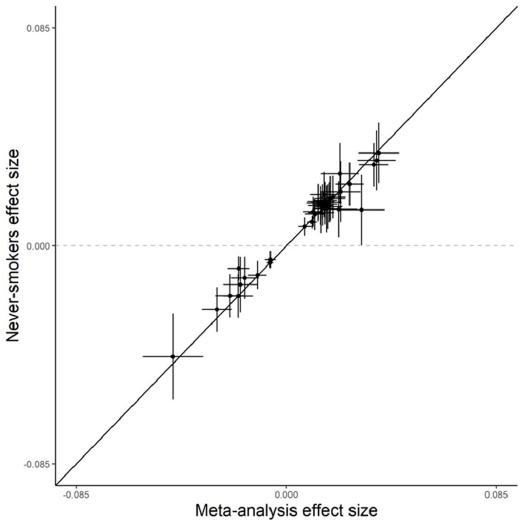
Comparison of meta-analysis effect size estimates and their 95 % confidence intervals in all participants (x-axis) and never-smokers (y-axis) for the 36 top CpG sites. All effect size estimates are adjusted for offspring sex, body mass index, smoking status, population stratification and technical covariates.

We then examined the dose-response relationship between maternal smoking and DNA methylation in the offspring. Methylation differences between the exposed and unexposed offspring became larger with increased smoking intensity across most CpG sites, e.g. each additional 3 cigarettes smoked per day during pregnancy was associated with 0.23 standard deviation (SD) increase in methylation level in cg05549655 in *CYP1A1* gene (Table 2). Figure 3 shows the visual representations of the dose-response effect of maternal smoking on offspring DNA methylation of top CpGs in four top loci.

**Table 2.** Association results for the leading CpG sites from each locus selected for the sensitivity and downstream analyses.

|  |  |  | Meta-analysis |  |  | Sensitivity and downstream analyses |  |  |  |  |  |  |  |  |
| --- | --- | --- | --- | --- | --- | --- | --- | --- | --- | --- | --- | --- | --- | --- |
|  |  |  | (5 studies)<br>N = 2821 |  |  | Never-smokers<br>N = 1298 |  |  |  | Paternal smoking<br>N = 1774 |  | Dose-response analysis<br>N = 1134 |  |  |
| CpG | Chr | Gene | β | SE | P-value | β | SE | P-value | β | SE | P-value | β | SE | P-value |
| cg15325070 | 1 |  | 0.014 | 0.003 | 9.23E-08 | 0.009 | 0.004 | 1.92E-02 | 0.000 | 0.003 | 9.16E-01 | 0.168 | 0.034 | 6.81E-07 |
| cg25189904 | 1 | GNG12 | -0.020 | 0.003 | 1.28E-10 | -0.018 | 0.004 | 1.87E-05 | -0.004 | 0.003 | 1.52E-01 | -0.146 | 0.032 | 5.85E-06 |
| cg14179389 | 1 | GFI1 | -0.028 | 0.003 | 5.00E-20 | -0.027 | 0.004 | 1.07E-09 | -0.004 | 0.003 | 1.77E-01 | -0.214 | 0.031 | 4.92E-12 |
| cg04749740 | 2 |  | 0.015 | 0.003 | 9.21E-08 | 0.015 | 0.004 | 9.64E-04 | 0.004 | 0.003 | 1.03E-01 | 0.061 | 0.034 | 6.98E-02 |
| cg06635952 | 2 | ANXA4 | 0.011 | 0.002 | 5.54E-08 | 0.011 | 0.003 | 1.88E-04 | 0.001 | 0.002 | 9.53E-01 | 0.073 | 0.029 | 1.12E-02 |
| cg11025974 | 2 | CACNB4 | 0.016 | 0.002 | 9.06E-11 | 0.018 | 0.004 | 1.57E-07 | -0.001 | 0.002 | 6.41E-01 | 0.169 | 0.033 | 2.19E-07 |
| cg14157435 | 2 | NRP2 | -0.046 | 0.006 | 1.15E-13 | -0.042 | 0.008 | 6.01E-07 | -0.014 | 0.006 | 2.80E-02 | -0.201 | 0.033 | 9.33E-10 |
| cg05783384 | 2 |  | 0.021 | 0.004 | 2.59E-08 | 0.014 | 0.005 | 1.15E-02 | 0.003 | 0.004 | 3.92E-01 | 0.097 | 0.033 | 3.26E-03 |
| cg05204104 | 2 | ARL4C | 0.017 | 0.003 | 4.81E-12 | 0.016 | 0.004 | 6.02E-05 | 0.006 | 0.003 | 7.96E-02 | 0.036 | 0.033 | 2.73E-01 |
| cg00174179 | 3 | RHOA | -0.006 | 0.001 | 7.01E-09 | -0.006 | 0.002 | 3.22E-04 | -0.002 | 0.001 | 5.21E-02 | -0.104 | 0.029 | 3.89E-04 |
| cg16449012 | 4 | FAM184B | 0.014 | 0.002 | 3.10E-08 | 0.021 | 0.004 | 4.42E-08 | 0.003 | 0.002 | 1.62E-01 | 0.119 | 0.03 | 7.81E-05 |
| cg05575921 | 5 | AHRR | -0.019 | 0.002 | 3.53E-18 | -0.011 | 0.003 | 1.19E-04 | -0.008 | 0.003 | 2.75E-03 | -0.134 | 0.027 | 5.91E-07 |
| cg21253335 | 5 |  | 0.017 | 0.003 | 1.25E-08 | 0.014 | 0.004 | 1.12E-03 | 0.004 | 0.003 | 1.17E-01 | 0.108 | 0.033 | 1.04E-03 |
| cg01952185 | 5 |  | 0.019 | 0.003 | 3.31E-12 | 0.017 | 0.004 | 5.81E-05 | 0.005 | 0.003 | 4.49E-02 | 0.128 | 0.03 | 1.67E-05 |
| cg05634495 | 6 |  | 0.016 | 0.003 | 3.14E-08 | 0.014 | 0.005 | 2.54E-03 | 0.002 | 0.003 | 5.00E-01 | 0.088 | 0.033 | 6.86E-03 |
| cg00794911 | 6 |  | -0.012 | 0.002 | 5.20E-10 | -0.007 | 0.003 | 1.25E-02 | -0.005 | 0.002 | 1.15E-02 | -0.115 | 0.034 | 6.42E-04 |
| cg25879142 | 7 |  | 0.018 | 0.003 | 1.79E-08 | 0.013 | 0.004 | 3.05E-03 | 0.003 | 0.003 | 3.73E-01 | 0.121 | 0.031 | 1.23E-04 |
| cg20117519 | 7 |  | 0.022 | 0.004 | 2.39E-08 | 0.022 | 0.006 | 1.70E-04 | 0.007 | 0.004 | 4.08E-02 | 0.112 | 0.033 | 6.72E-04 |
| cg19089201 | 7 | *MYO1G* | 0.035 | 0.003 | 1.18E-31 | 0.03 | 0.004 | 5.75E-12 | 0.007 | 0.003 | 1.51E-02 | 0.225 | 0.032 | 2.24E-12 |
| cg04598670 | 7 |  | -0.019 | 0.003 | 8.33E-08 | -0.016 | 0.005 | 4.54E-03 | -0.007 | 0.003 | 3.95E-02 | -0.178 | 0.034 | 1.54E-07 |
| cg25949550 | 7 | *CNTNAP2* | -0.007 | 0.001 | 5.41E-27 | -0.007 | 0.001 | 1.45E-14 | -0.002 | 0.001 | 6.57E-03 | -0.157 | 0.022 | 1.65E-12 |
| cg11207515 | 7 | *CNTNAP2* | -0.023 | 0.003 | 1.03E-13 | -0.02 | 0.004 | 2.75E-06 | -0.011 | 0.003 | 2.46E-04 | -0.145 | 0.032 | 7.02E-06 |
| cg15578140 | 7 | *MIR548F3* | 0.011 | 0.002 | 7.54E-09 | 0.014 | 0.003 | 7.89E-06 | -0.001 | 0.002 | 5.70E-01 | 0.152 | 0.031 | 1.12E-06 |
| cg17199018 | 8 | *ZNF395* | -0.017 | 0.003 | 3.85E-09 | -0.016 | 0.004 | 7.10E-05 | -0.003 | 0.003 | 3.10E-01 | -0.121 | 0.033 | 2.63E-04 |
| cg14563637 | 9 |  | 0.016 | 0.003 | 1.19E-09 | 0.016 | 0.004 | 9.26E-05 | 0.003 | 0.002 | 1.69E-01 | 0.092 | 0.031 | 3.26E-03 |
| cg14540913 | 9 | *PRRX2* | 0.013 | 0.002 | 7.57E-09 | 0.015 | 0.003 | 1.81E-05 | 0.001 | 0.002 | 5.79E-01 | 0.148 | 0.032 | 2.62E-06 |
| cg13822849 | 9 | *OLFM1* | 0.007 | 0.001 | 2.87E-08 | 0.007 | 0.002 | 5.84E-05 | 0.003 | 0.002 | 2.90E-02 | 0.126 | 0.03 | 3.24E-05 |
| cg11813497 | 10 | *FRMD4A* | 0.026 | 0.003 | 9.27E-19 | 0.022 | 0.004 | 1.46E-07 | -0.001 | 0.003 | 6.26E-01 | 0.117 | 0.032 | 2.81E-04 |
| cg05697249 | 11 | *C11orf52* | 0.015 | 0.002 | 1.30E-11 | 0.015 | 0.003 | 8.68E-06 | 0.003 | 0.002 | 1.54E-01 | 0.136 | 0.035 | 8.25E-05 |
| cg18493761 | 11 |  | 0.037 | 0.004 | 7.30E-21 | 0.032 | 0.006 | 4.99E-08 | 0.008 | 0.004 | 2.60E-02 | 0.154 | 0.034 | 4.82E-06 |
| cg05549655 | 15 | *CYP1A1* | 0.010 | 0.001 | 1.04E-28 | 0.009 | 0.001 | 6.03E-13 | 0.002 | 0.001 | 4.28E-02 | 0.23 | 0.028 | 1.92E-16 |
| cg13834112 | 15 |  | 0.013 | 0.002 | 3.55E-08 | 0.018 | 0.004 | 6.39E-07 | 0.001 | 0.002 | 8.10E-01 | 0.12 | 0.031 | 1.24E-04 |
| cg00253658 | 16 |  | 0.037 | 0.004 | 3.44E-19 | 0.032 | 0.006 | 6.54E-08 | 0.005 | 0.005 | 2.81E-01 | 0.166 | 0.033 | 4.83E-07 |
| cg04358214 | 16 | *C16orf70* | 0.022 | 0.004 | 9.56E-08 | 0.021 | 0.006 | 5.93E-04 | 0.005 | 0.004 | 2.42E-01 | 0.09 | 0.031 | 3.94E-03 |
| cg12984635 | 19 | *ETHE1* | 0.015 | 0.002 | 2.17E-09 | 0.014 | 0.003 | 5.87E-05 | 0.001 | 0.002 | 7.33E-01 | 0.116 | 0.031 | 1.49E-04 |
| cg06758350 | 21 | *RUNX1* | 0.030 | 0.005 | 1.68E-10 | 0.016 | 0.007 | 2.03E-02 | 0.002 | 0.005 | 7.09E-01 | 0.14 | 0.034 | 3.92E-05 |

The analyses were conducted separately in each participating cohort, adjusted for offspring age, sex, body mass index, smoking status and genetic ancestry, and combined using inverse-variance weighted fixed effects meta-analysis. Chr = chromosome; $\beta$ = effect size estimate, SE = standard error.

To assess potential unmeasured confounding and to establish a causal intra-uterine effect between maternal smoking and the offspring DNA methylation we used paternal smoking as a negative control. Maternal smoking and paternal smoking showed similar directions of effect; however, the effect estimates for exposure to paternal smoking were considerably smaller (Table 2). Adjusting for paternal smoking had no significant effect on maternal smoking estimates (**Supplementary Table 4**).

**Figure 3.**
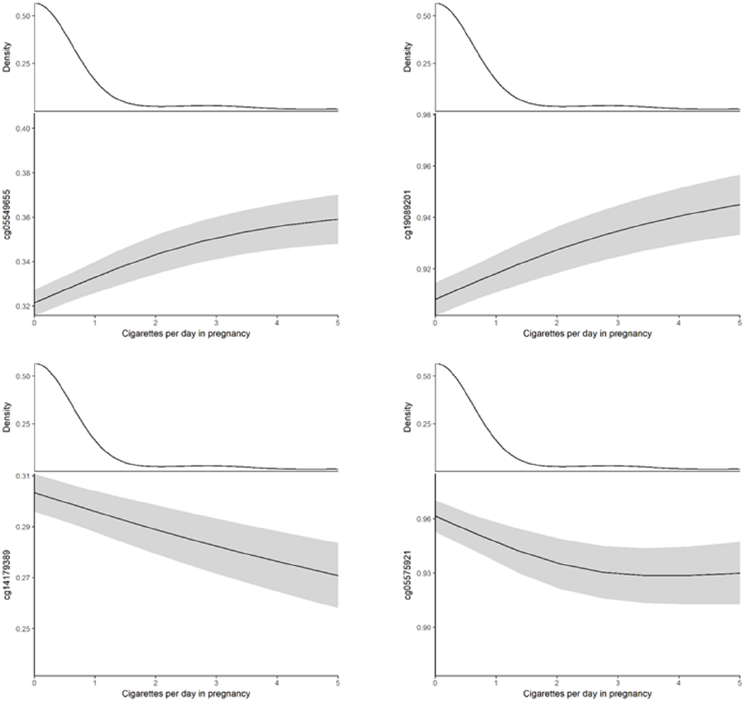
Visualization of the dose-response effect of maternal smoking on offspring DNA methylation for top 4 CpG sites in four gene regions (AHRR, CYP1A1, MYO1G, GFI1). Prediction estimates and their 95 % confidence intervals plotted based on generalized additive mixed models, with other covariates (offspring sex, body mass index, smoking status, population stratification and technical covariates) set at their mean (continuous variables) or mode (categorical variables). The plots are truncated at 5 cigarettes per day in pregnancy (containing 94 % of full data).

We performed a longitudinal analysis to examine whether the maternal smoking associated alterations in DNA methylation persisted from early adulthood (age 30-31y) into midlife (age 46-48y) in the NFBC 1966 and ALSPAC mothers cohorts. We found no evidence for change in direction or magnitude of associations in DNA methylation between the two time points (Figure 4), suggesting that DNA methylation levels remains relatively stable for several decades after prenatal exposure to maternal smoking.

**Figure 4.**
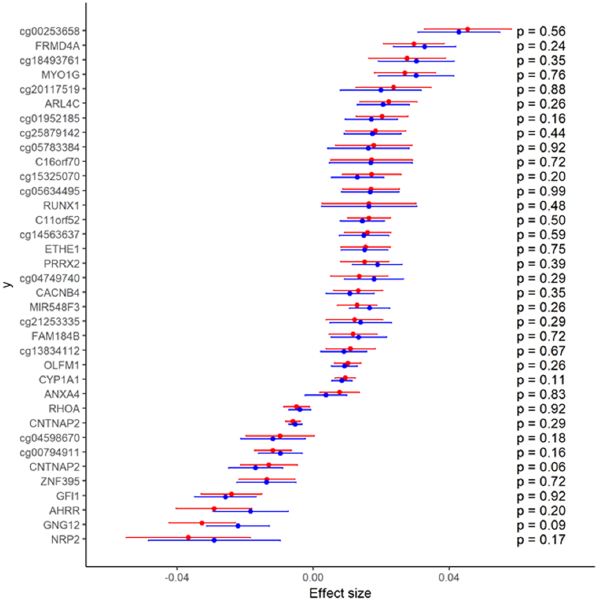
Longitudinal analysis of association between exposure to maternal smoking and offspring DNA methylation. Effect estimates (adjusted for offspring sex, body mass index, smoking status, population stratification and technical covariates) and their 95 % confidence intervals at age 30-31 years (blue) and age 46-48 years (red) for top CpG sites and P values for the equality of the effect estimates.

### Mendelian randomization analysis

We estimated the causal effects of DNA methylation changes on disease outcomes using MR. We extracted the effect sizes of SNP-CpG associations for the 69 differentially methylated CpGs available in the Accessible Resource for Integrated Epigenomic Studies (ARIES) mQTL database^16^(http://www.mqtldb.org/), and found strong instruments for 15 CpG sites. Of these 15 CpG sites, three (cg15578140 in microRNA 548f-3 (*MIR548F3*), cg09935388 in Growth Factor Independent Protein 1 (*GFI1*), cg04598670 (unknown gene)) showed potential causal associations with inflammatory bowel diseases and one (cg25189904 in Guanine Nucleotide Binding Protein Gamma 12 (*GNG12*)) with Schizophrenia (*P*_*FDR*_ < 0.05, Table 3).

**Table 3.** Mendelian randomization analysis of top differentially methylated CpGs tested against 106 diseases.

| exposure | gene | disease | $\beta$ | se | pval | pval FDR | unit |
| --- | --- | --- | --- | --- | --- | --- | --- |
| cg15578140 | *MIR548F3* | Inflammatory bowel disease | -0.104 | 0.018 | 3.73E-09 | 2.54E-07 | log odds |
| cg09935388 | *GFI1* | Inflammatory bowel disease | -0.152 | 0.034 | 7.27E-06 | 0.000218 | log odds |
| cg04598670 | unknown | Inflammatory bowel disease | -0.410 | 0.091 | 7.27E-06 | 0.000524 | log odds |
| cg09935388 | *GFI1* | Crohn's disease | -0.162 | 0.040 | 4.74E-05 | 0.000712 | log odds |
| cg04598670 | unknown | Crohn's disease | -0.439 | 0.108 | 4.74E-05 | 0.001708 | log odds |
| cg09935388 | *GFI1* | Ulcerative colitis | -0.160 | 0.042 | 0.000147 | 0.001467 | log odds |
| cg04598670 | unknown | Ulcerative colitis | -0.433 | 0.114 | 0.000147 | 0.003521 | log odds |
| cg25189904 | *GNG12* | Schizophrenia | -0.222 | 0.053 | 3.37E-05 | 0.001819 | log odds |

Data are given as beta coefficients and their standard errors. Only significant associations are shown. *GFI1* = Growth Factor Independent Protein 1; *MIR548F3* = microRNA 548f-3; *GNG12* = Guanine Nucleotide Binding Protein Gamma 12. FDR = false discovery rate.

### Mediation analysis

We then sought to test whether methylation changes in these four CpGs mediated the association between maternal smoking and disease outcomes. However, since the prevalence of inflammatory bowel disease is relatively low in general population, we assessed the associations of maternal smoking and CpGs on irritable bowel syndrome (IBS), which is a constellation of functional gastrointestinal disorder symptoms. These data were obtained from self-administered questionnaires in NFBC1966 at 46 years^17^. Prevalence of schizophrenia is also low in the general population. Therefore, instead of diagnosed schizophrenia, we used personality trait scales measuring schizotypal and affective symptoms as an outcome. Such personality scales were derived from questionnaires available in the NFBC 1966 data at 31 years and they can be used to identify subjects with latent personality with genetic vulnerability for schizophrenia^18^. We found evidence for cg25189904 mediating the association between exposure to maternal smoking and Bipolar II Scale (*P* = 0.024) and Hypomanic Personality Scale (*P* = 0.018) (Figure 5 A and Figure 5 B). The estimated mediated proportions were 30% and 28%, respectively (**Supplementary Table 5**).

**Figure 5.**
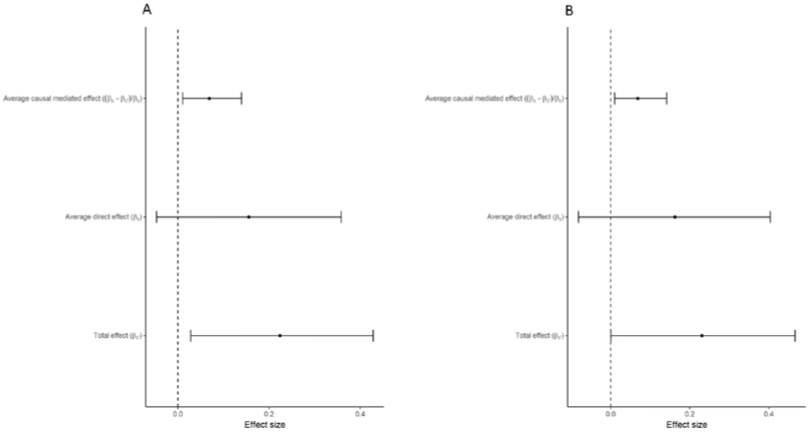
Mediation analysis examining the indirect effect of maternal smoking during pregnancy on Bipolar II Scale (A) and Hypomanic personality scale (B) through differential methylation of cg25189904 in GNG12. Data are shown as beta estimate for effect size and 95 % CI.

## Discussion

We combined data from five studies in adolescents and adults to examine the association between maternal smoking during pregnancy and DNA methylation in the offspring from 16 until 48 years of age. We identified 69 differentially methylated CpGs in 36 genomic regions. The top differentially methylated CpG sites showed a clear dose-response relationship with number of cigarettes smoked during pregnancy. The associations observed in adulthood were robust to adjustment for multiple potential confounding factors and persisted into middle age with no significant change in direction and magnitude of associations. Mendelian randomization and mediation analyses suggested that alterations in DNA methylation may link maternal smoking during pregnancy to increased risk of psychiatric morbidity and potentially with inflammatory bowel disease in the exposed offspring.

The findings of our study confirm and extend the results of earlier reports by demonstrating that maternal smoking during pregnancy is associated with alterations in offspring DNA methylation not only in newborns^11,19,20^, children and adolescents^12,13^, but also in adults, several decades following the exposure. The similarity in differentially methylated CpG sites and the consistency in direction of methylation changes between our study and earlier EWAS imply that the smoke-exposure induced methylation changes may be soma-wide and persist throughout life. However, the effects of smoking may also be targeted to specific regions of the epigenome, as indicated by the observations that both prenatal smoke exposure and active smoking affect the methylation patterns of same gene regions, e.g. *AHRR* and *CYP1A1*, which are involved in chemical detoxification^10^. Because of these similar effects, the methylation changes found in people exposed to prenatal smoking may also reflect current or past smoking by the people themselves or some other passive smoking exposure. Adjusting for offspring active smoking did not substantially change the results in the present study. However, parental smoking is known to associate with their offspring’s smoking behavior also via genetic predisposition^21,22^and thus own smoking may serve as a mediator on the path between maternal smoking and DNA methylation. Therefore, simply adjusting for own smoking can lead to erroneous conclusions about the direct effects of maternal smoking^23^. We therefore performed a sensitivity analysis including only offspring who themselves had never smoked in their life and found that the associations were similar across all CpG sites as in the full meta-analysis.

We also used paternal smoking as a negative control by comparing the associations of maternal smoking during pregnancy and paternal smoking with offspring methylation and found that the effect estimates were substantially greater for maternal smoking, and adjusting for paternal smoking had virtually no effect on maternal smoking estimates. This indicates it is unlikely that the associations between maternal smoking and offspring methylation were attributable to post-natal passive smoking exposure or some unmeasured confounding. These results together with the finding of a clear dose-dependent relationship of methylation with increased smoking intensity during pregnancy suggest a direct biological effect of in utero exposure to cigarette smoke on DNA methylation.

The longitudinal analysis showed that differentially methylated CpGs observed around age 30 persisted into middle age (around age 48) without significant change in direction or magnitude of methylation levels. This corroborates the findings of recent smaller studies, which found several differently methylated CpGs in middle-aged women exposed to maternal smoking in utero^14,15^and suggests that some of the prenatal smoking exposure associated methylation changes are largely irreversible and unaffected by age and/or environmental exposures later in life. To assess whether such persistent changes in DNA methylation are causally implicated with disease, we performed a Mendelian randomization analysis using summary data from large epigenome-wide association studies^24^. We found evidence for potential causal associations for three CpGs (cg15578140, cg09935388, cg04598670) with inflammatory bowel disease and one CpG (cg25189904) with Schizophrenia. To confirm these potential causal associations, we also performed a formal mediation analysis in the NFBC 1966 cohort and found evidence for differential methylation in cg25189904 mediating the association between maternal smoking and Bipolar II Scale and Hypomanic Personality Scale, explaining 30% and 28% of the total effect, respectively. These results corroborate the findings of previous observational studies, that maternal smoking during pregnancy is associated with increased risk of psychiatric morbidity in the exposed offspring^25–28^. However, we found no evidence for mediating effect of differential methylation cg15578140, cg09935388 and cg04598670 in the association of maternal smoking and irritable bowel syndrome. Such discrepant results could be due to relatively small sample size in the mediation analysis, or because the irritable bowel syndrome is not a good proxy for inflammatory bowel disease, or because the causal effect estimates for inflammatory bowel disease in the MR analysis were biased due to, for example, pleiotropic effects of genetic instruments on the outcome. Thus, additional studies are needed to assess whether prenatal smoking is associated with increased risk of inflammatory bowel disease in the exposed offspring and whether alterations in DNA methylation mediates this association.

Our results provides a novel potential biomarker for psychiatric research. Experimental studies suggest that *GNG12* is an important regulator of inflammatory signaling in microglia cells, which are the resident macrophages of the central nervous system^29^. A role of inflammation in the etiology of schizophrenia and psychotic illness has been suggested^30,31^, and in line with this, a large meta-analysis of 2424 cases and over 1.2 million controls indicated that childhood central nervous system infections are associated with nearly two-fold risk of schizophrenia in adulthood^32^. Whereas the pathogenic processes for psychiatric disorders most likely occur in the brain, studies have shown that factors affecting the brain processes can leave a methylation signature in the blood that mirror the methylation sites in the brain^33^. Such mirror sites can occur if the exposure occurs during early stages of prenatal development, thus affecting both the brain and peripheral tissues^33^. Further studies are needed to validate our findings and investigate the biological relevance of *GNG12* in the primary disease tissue.

Our study has both strengths and limitations. The large sample size of males and females and similar ages from different cohorts enabled us to obtain precise estimate of the long-term effects of maternal smoking on DNA methylation. Several downstream analyses and use of paternal smoking as a negative control allowed us to distinguish the associations from potential confounding, and the follow-up analysis from young adulthood to middle-age allowed us to examine the persistency of methylation changes. The limitations are that maternal smoking was determined from self-reported questionnaires. As self-reports may be biased by under-reporting or recall bias, our findings may be underestimate true effects. In the ALSPAC mothers cohort the adult offspring reported their mothers’ smoking, although this could also be subject to recall bias. False reporting may also concern the adolescents in our study since they might have been reluctant to disclose their true smoking behavior, although in the IOWBC adolescent smoking was confirmed by urinary cotinine measurement. Another limitation is that the subjects in the ALSPAC children and ALSPAC mothers cohorts are related individuals. However, excluding either one of the related ALSPAC data sets did not notably affect the results (data not shown).

## Conclusions

Maternal smoking during pregnancy has long-lasting effects on offspring epigenome. DNA methylation may represent a biological mechanism through which maternal smoking is associated with increased risk of psychiatric morbidity and potentially inflammatory bowel disease in the exposed offspring.

## Methods

### Study cohorts

#### Northern Finland Birth Cohort 1966

The Northern Finland Birth Cohort 1966 previously described in detail^34,35^, targeted all pregnant women, residing in the two northernmost provinces of Finland with expected dates of delivery between 1 January and 31 December 1966. Over 96 % of eligible women participated in the study, thus comprising 12,055 mothers followed prospectively on average from 16^th^ gestational week and 12,058 live born children. In 1997, at offspring age of 31 years, all cohort participants with known addresses were sent a postal questionnaire on health and lifestyle and those living in Northern Finland or Helsinki area were invited to a clinical examination which included blood sampling. In total, both questionnaire and clinical data were collected for 6,007 participants. DNA was successfully extracted for 5,753 participants from fasted blood samples^36^. In 2012, all individuals with known address in Finland were sent postal questionnaires and an invitation for clinical examination. Both questionnaire and clinical data was collected for 5,539 participants. DNA methylation at 31 years was extracted for 807 randomly selected subjects of whom both questionnaire and clinical data with cardio-metabolic measures were available at both 31 and 46 years. Of these individuals, DNA methylation data at 46 years was extracted for 766 subjects.

#### Northern Finland Birth Cohort 1986

The Northern Finland Birth Cohort 1986 includes all mothers (prospective data collection from 10^th^ gestational week) with children whose expected date of delivery fell between July, 1st 1985 and June, 30th 1986, in the two northernmost provinces of Finland (99% of all births during that time) ^37^. The cohort consists of 9,362 women and 9,432 live-born children. In 2001, all individuals with known address received a postal questionnaire on health and lifestyle and invitation to a clinical examination. DNA were extracted from fasting blood samples and DNA methylation were measured for 442 randomly selected subjects with full data available.

In both NFBC cohorts, complete data included singleton births and subjects with complete set clinical follow-up and DNA methylation data; excluding subjects with missing information and twins. A written informed consent for the use of the data including DNA was obtained from all study participants and their parents. Ethical approval for the study was received from Ethical Committee of Northern Osthrobothnia Hospital District and Oulu University, Faculty of Medicine.

#### Isle of Wight Birth Cohort

Isle of Wight Birth cohort is a general population based birth cohort recruited on the Isle of Wight in 1989 to assess the role of heredity and environment on development of allergic disorders and allergen sensitization. The details of this birth cohort have been described in previous reports^38^. In brief, both the Isle of Wight and the study population are 99% Caucasian. Ethics approvals were obtained from the Isle of Wight Local Research Ethics Committee (now named the National Research Ethics Service, NRES Committee South Central –Southampton B) at recruitment and for the 1, 2, 4, 10 and 18 years follow-up. Exact age at 18-year follow-up was calculated from the date of blood sample collection for the 18-year follow-up and the date of birth. DNA methylation in peripheral blood samples was analyzed from randomly selected subjects (n = 257) at the 18-year follow-up.

#### Avon Longitudinal Study of Parents and Children

Pregnant women resident in the former county of Avon, UK with expected dates of delivery 1st April 1991 to 31st December 1992 were invited to take part in the study. The initial number of pregnancies enrolled is 14,541 (for these at least one questionnaire has been returned or a “Children in Focus” clinic had been attended by 19/07/99). Of these initial pregnancies, there was a total of 14,676 foetuses, resulting in 14,062 live births and 13,988 children who were alive at 1 year of age^39,40^.

The Accessible Resource for Integrated Epigenomic Studies (ARIES) is a sub study of ALSPAC, which includes 1,018 mothers and their children for whom methylation data has been created^41^. The ARIES participants were selected based on the availability of DNA samples at two time points for the women (antenatal [mean age 30 years] and at follow-up [mean age 48 years] when the offspring were adolescents) and three time points for their offspring (neonatal, childhood [mean age 7.5 years], and adolescence [mean age 17.1 years]). A web portal allows openly accessible browsing of aggregate ARIES DNA methylation data (ARIES-Explorer) (http://www.ariesepigenomics.org.uk/). Please note that the study website contains details of all the data that is available through a fully searchable data dictionary and variable search tool: http://www.bristol.ac.uk/alspac/researchers/our-data/. Ethical approval for the study was obtained from the ALSPAC Ethics and Law Committee and the Local Research Ethics Committees.

### Definition of maternal smoking during pregnancy

In NFBCs and ALSPAC studies, expectant mothers were asked whether they had smoked cigarettes before or at the beginning of the pregnancy, how many years they had smoked, the number of cigarettes smoked per day, and whether they had changed their smoking habits during the pregnancy. Offspring were considered to be prenatally exposed to cigarette smoking if mother reported smoking regularly (at least one cigarette per day) from pregnancy week 8 onwards. The ALSPAC mothers were also asked whether their mothers had smoked, and were asked whether they had smoked when they were pregnant with them. In the IOWBC, maternal smoking status in pregnancy was self-reported and defined as any smoking in pregnancy or no smoking during pregnancy.

### Measurement of DNA methylation

Methylation of genomic DNA was quantified using the Illumina HumanMethylation450 array (ALSPAC, ARIES, IOWBC, and NFBC1966 at age 31, NFBC1986) or Illumina EPIC array (NFBC1966 at age 46) according to manufacturer’s instructions. Bisulfite conversion of genomic DNA was performed using the EZ DNA methylation kit according to manufacturer’s instructions (Zymo Research, Orange, CA).

### Quality control of methylation data

In NFBCs and IOWBC quality control and quantile normalization for DNA methylation data was adapted from the CPACOR pipeline ^42^. Illumina Background Correction was applied to the intensity values, a detection p-value threshold was set at *P* < 10^−16^, and samples with call rate < 98 % were excluded. Quantile normalization was done separately for six probe-type categories, and these normalized intensity values were used to calculate the methylation beta value at each CpG site, ranging between 0 (no methylation) and 1 (full methylation). Probes with call rate < 95 % were excluded from the analyses. A principal component analysis (PCA) was carried out for array control probes, and the first 30 principal components (PCs) were used as explanatory variables in the subsequent regression models. White blood cell subpopulation estimates were obtained using the software provided by Houseman et al. ^43^ and these estimates were also added as covariates in the regression models. In ARIES, the DNA methylation wet-laboratory and pre-processing analyses were performed as previously described^41^. In brief, samples from all time points were distributed across slides using a semi-random approach to minimize the possibility of confounding by batch effects. Samples failing quality control (average probe *P* value ≥0.01, those with sex or genotype mismatches) were excluded from further analysis and scheduled for repeat assay, and probes that contained <95% of signals detectable above background signal (detection *P* value <0.01) were excluded from analysis. Methylation data were pre-processed using R software, with background correction and subset quantile normalization performed using the pipeline described by Touleimat and Tost ^44^.

## Statistical analyses

### Meta-analysis of 6073 CpG sites in five studies

Study design and analytical flow of the study are shown in Figure 1 and the data availability for each analysis is presented in Table 4. All analyses were conducted using R software ^45^. Linear regression was used to examine the association between sustained maternal smoking during pregnancy (from pregnancy week 8 onwards) and offspring peripheral blood DNA methylation at 6073 CpG sites that were previously identified to be differentially methylated in newborns exposed to maternal smoking in utero in recent epigenome-wide association study (EWAS) (false discovery rate corrected *P* value < 0.05). The final model was adjusted for study specific covariates as necessary (offspring’s sex, age, BMI, smoking status, and maternal age and social class as well as genetic PCs to account for population stratification). The model was run independently in each study, and the results were then meta-analyzed over all five studies (NFBC1986, NFBC1966 (age 16y & 31y), IOWBC (age 18y) and ALSPAC mothers (age 30y), ALSPAC children (age 17y). Statistical significance level was set at *P* < 1×10^−7^, which corresponds approximately to a Bonferroni-corrected significance level of 0.05 for 450,000 independent tests. Such a conservative threshold was robust and thus the significant probes were considered worthy of further examination in a series of sensitivity and downstream analyses. The leading CpG site from each gene region (1 Mb window centered on the CpG site with the strongest association) was selected for these analyses.

**Table 4.** Data available in each study for different analyses.

| Study | Mean age at methylation data collection (years) | Association results for cord-blood-methylation associated CpG sites | Sensitivity and downstream analyses for top CpG sites |  |  |  |
| --- | --- | --- | --- | --- | --- | --- |
|  |  |  | Never-smokers | Paternal smoking | Amount of cigarettes smoked in pregnancy | Longitudinal methylation data (mean age in years at 2<sup>nd</sup> time point) |
| NFBC1986 | 16 | Yes | Yes | Yes | Yes | No |
| NFBC1966 | 31 | Yes | Yes | Yes | Yes | Yes (47) |
| ALSPAC mothers | 30 | Yes | Yes | No | No | Yes (48) |
| ALSPAC children | 17 | Yes | Yes | Yes | No | No |
| IOWBC | 18 | Yes | Yes | No | No | No |

## Sensitivity analyses

### Impact of offsprings’ own smoking on their DNA methylation

To assess the impact of participants’ own smoking on methylation level by maternal smoking exposure, the same regression model was run excluding all participants who reported smoking regularly, defined in NFBC1966 as smoking at least one cigarette per day for one year or more during their life. In the ALSPAC mothers’ cohort smoking behavior was queried at two time points. At age 30y, women were asked whether they had smoked regularly before pregnancy. At age 48y, women were asked whether they were current or former smokers, and in case of the latter, whether they had smoked every day. From these data a dichotomous variable for any smoking for each of the time points was derived. In the IOWBC, participant’s own smoking status was defined as having ever or never smoked asked via questionnaire administered at age 18y.

### Impact of mother’s smoking intensity on offspring DNA methylation

Further analyses were performed to investigate whether intensity of maternal smoking during pregnancy had differential impact on the level of offspring DNA methylation. For this, the association between the number of cigarettes smoked per day during pregnancy and offspring DNA methylation was assessed in the NFBC studies.

### Negative control design to distinguish intrauterine effects from confounding

Potential unmeasured confounding was examined in the NFBC studies by using paternal smoking status during pregnancy as a negative control. This method compares the associations of maternal and paternal smoking during pregnancy with offspring methylation outcomes. Use of paternal smoking as a negative control is based on the assumption that the biological effects of paternal smoking on intrauterine exposure are negligible compared to the effects of maternal smoking during pregnancy. If there is an intrauterine effect of cigarette smoke exposure, the associations are expected to be stronger for maternal smoking than paternal smoking behavior. If effects are of similar magnitude, the associations between maternal smoking during pregnancy and offspring methylation are likely attributable to unmeasured confounding, either by shared environmental or genetic factors^46^.

### Persistence of DNA methylation into adulthood

We also examined whether the methylation changes associated with maternal smoking persisted into middle-age. DNA methylation data were available at two time-points in NFBC 1966 (age 31y and 46y) and ALSPAC mother (age 30y and 48y). Generalized least squares were used to examine the longitudinal change in association between exposure to maternal smoking and DNA methylation. Time point of measurement and its interaction with the exposure were added as additional terms to the regression model, and the model residuals were allowed to be correlated within each individual and be heteroskedastic between time points. The effect estimates at both time points can be derived from this model, and the test for equality of the estimates at both time points is equivalent to testing the interaction term being equal to zero ^47^. All the above-described analyses were conducted separately in the studies where corresponding data were available (Table 4), and meta-analyzed using an inverse-variance weighted fixed-effects model.

### Mendelian randomization analysis for the effect of DNA methylation on disease outcomes

We next sought to assess the potential causal relationship between DNA methylation as the exposure and 106 different diseases as outcomes available through the MR-Base platform (available at http://www.mrbase.org/) using two-sample Mendelian randomization (MR). The two-sample MR approach uses gene-exposure and gene-outcome associations from different data sources of comparable populations and allows the interrogation of summary estimates available from large genome-wide association (GWAS) consortia^24^. If instrumental variable assumptions for the genes associated with the proxies are fulfilled^48^, then MR estimates can give evidence for a causal effect of exposure on the outcome.

We first looked up proxy single nucleotide polymorphisms (SNPs) for each of the 69 top maternal-smoking associated CpG sites in the publicly available ARIES database containing methylation quantitative trait loci (mQTL) at four different life stages (birth, childhood, adolescence, middle age) in human blood^41^. We selected SNPs associated with each CpG at *P* < 10^−7^ at any of the other four time points. After clumping SNPs (using 1 MB window and *R*^2^ < 0.001) and pruning the CpG sites to one per locus, we found strong instruments for 15 CpG sites (**Supplementary Table 6**). These SNP-CpG associations were consistent across all time points (**Supplementary Figure 1**), except rs4306016-cg01825213 association, which was excluded from the final MR analysis. We selected the SNP-CpG and SNP-disease effects sizes at middle age and aligned these to the same allele. MR effect estimates were then calculated using Wald ratio or, in case of cg04598670, which had two SNP instruments available, inverse variance weighted method. The resulting effect estimate represents the change in outcome per unit increase in the exposure.

### Mediation analysis

The CpGs that showed evidence for causal relationship with disease outcomes in the MR analysis were tested for mediation in the association between maternal smoking during pregnancy and disease outcomes using the NFBC1966 data at 31y and 46y. We performed model-based causal mediation analysis using R package ‘mediation’^49^by first estimating both the effect of maternal smoking on the CpG site and the effect of CpG site on the outcome, adjusted for exposure to maternal smoking (Figure 6). Both of these effects were additionally adjusted for sex, offspring’s own smoking and technical covariates. We generated the estimates for the total effect, average direct effect and average causal mediation effect using quasi-Bayesian Monte Carlo method based on normal approximation with 2000 simulations, with robust standard errors. The proportion that the mediating CpG explains of the association between maternal smoking and disease outcome was calculated as described^50^.

**Figure 6.**
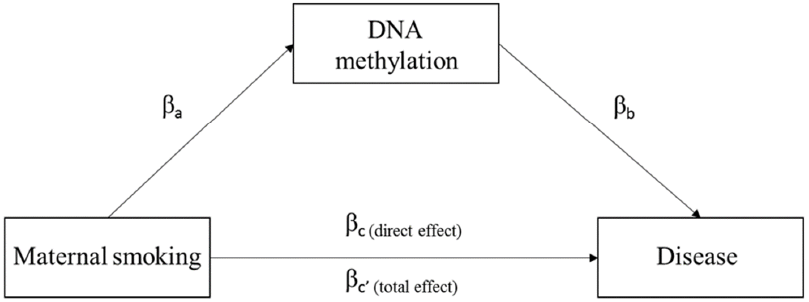
A mediation model for the association between maternal smoking and offspring disease outcomes. βa represent the effect estimate for smoking on DNA methylation (CpG = maternal smoking + covariates); βb represents the effect estimate for CpG on disease (disease = CpG + covariates); βc represent the direct effect (no mediation) estimate for maternal smoking on disease (disease = maternal smoking + covariates; βc’ represents the total effect estimate on disease (disease = maternal smoking + covariates + CpG).

#### Competing interests: The authors declare no competing interests.

Funding/Support: This project was supported by the Academy of Finland EGEA-project (285547),Biocenter, University of Oulu, Finland (75617), NHLBI grant 5R01HL087679-02 through the STAMPEED program (1RL1MH083268-01), ERDF European Regional Development Fund Grant no. 539/2010 A31592, the EU H2020‐‐PHC-2014 DynaHEALTH action (grant agreements No. 633595), EU H2020-HCO-2004 iHEALTH Action (grant agreement 643774), EU H2020-PHC-2014 ALEC Action (grant agreement No. 633212), EU H2020-SC1-2016-2017 LifeCycle Action (grant agreement No 733206), EU H2020-MSCA-ITN-2016 CAPICE Action (grant agreement 721567) and MRC Grant nro MR/M013138/1.

The Isle of Wight Birth Cohort study has been supported by the National Institutes of Health USA (Grant no. R01 HL082925 (PI: Arshad), R01 AI091905 and R01 HL132321 (PI: Karmaus), and R01 AI121226 (MPI: Zhang and Holloway) and Asthma UK (Grant no. 364). JWH and FIR are supported by the Ageing Lungs in European Cohorts (ALEC) Study (www.alecstudy.org), which has been funded by the European Union’s Horizon 2020 Research and Innovation programme under grant agreement No. 633212.

Data contributions by the ALSPAC study were supported by the Integrative Epidemiology Unit, which receives funding from the UK Medical Research Council and the University of Bristol (MC_UU_12013_1 and MC_UU_12013_2). This work was also supported by CRUK (grant number C18281/A19169) and the ESRC (grant number ES/N000498/1). The UK Medical Research Council and the Wellcome Trust (Grant ref: 102215/2/13/2) and the University of Bristol provide core support for ALSPAC. The Accessible Resource for Integrated Epigenomics Studies (ARIES) was funded by the UK Biotechnology and Biological Sciences Research Council (BB/I025751/1 and BB/I025263/1). R.C.R is a de Pass VC Research Fellow at the University of Bristol. T.G.R is a UKRI Innovation Research Fellow (MR/S003886/1). GWAS data used to identify the mQTLs for the ALSPAC offspring was generated by Sample Logistics and Genotyping Facilities at Wellcome Sanger Institute and LabCorp (Laboratory Corporation of America) using support from 23andMe. Genotyping for the ALSPAC women was supported by the Wellcome Trust (grant reference WT088806). A comprehensive list of grants funding is available on the ALSPAC website (http://www.bristol.ac.uk/alspac/external/documents/grant-acknowledgements.pdf). The funders had no role in study design, data collection and analysis, decision to publish or preparation of the manuscript.

## Acknowledgements

We are extremely grateful to all the participants and families who took part in this study, the midwives for their help in recruiting them, and the whole research teams in each cohort, which includes interviewers, computer and laboratory technicians, clerical workers, research scientists, volunteers, managers, receptionists and nurses.

Author Contributions: Wiklund P, Karhunen V and Järvelin M-R had full access to all of the data in the study and take responsibility for the integrity of the data and the accuracy of the data analysis.

Concept and design: Wiklund P, Karhunen V, Rodriguez A, Relton C, and Järvelin M-R

Drafting of the work: Wiklund P, Karhunen V, Järvelin M-R

Critical revision of the manuscript for important intellectual content: All authors

Statistical analysis: Karhunen V., Richmond RC., Rezwan FI.

Data availability: the data that support the findings of this study are available are available from the corresponding author upon request.

## References

1 Anblagan, D. et al. Maternal smoking during pregnancy and fetal organ growth: a magnetic resonance imaging study. PloS one 8, e67223, doi:10.1371/journal.pone.0067223 (2013).

2 Horta, B. L., Victora, C. G., Menezes, A. M., Halpern, R. & Barros, F. C. Low birthweight, preterm births and intrauterine growth retardation in relation to maternal smoking. Paediatric and perinatal epidemiology 11, 140–151 (1997).

3 Shah, N. R. & Bracken, M. B. A systematic review and meta-analysis of prospective studies on the association between maternal cigarette smoking and preterm delivery. American journal of obstetrics and gynecology 182, 465–472 (2000).

4 Cupul-Uicab, L. A. et al. In utero exposure to maternal tobacco smoke and subsequent obesity, hypertension, and gestational diabetes among women in the MoBa cohort. Environmental health perspectives 120, 355–360, doi:10.1289/ehp.1103789 (2012).

5 Power, C., Atherton, K. & Thomas, C. Maternal smoking in pregnancy, adult adiposity and other risk factors for cardiovascular disease. Atherosclerosis 211, 643–648, doi:10.1016/j.atherosclerosis.2010.03.015 (2010).

6 Ng, S. P. & Zelikoff, J. T. Smoking during pregnancy: subsequent effects on offspring immune competence and disease vulnerability in later life. Reproductive toxicology 23, 428–437, doi:10.1016/j.reprotox.2006.11.008 (2007).

7 Doherty, S. P., Grabowski, J., Hoffman, C., Ng, S. P. & Zelikoff, J. T. Early life insult from cigarette smoke may be predictive of chronic diseases later in life. Biomarkers : biochemical indicators of exposure, response, and susceptibility to chemicals 14 Suppl 1, 97–101, doi:10.1080/13547500902965898 (2009).

8 Hofhuis, W., de Jongste, J. C. & Merkus, P. J. Adverse health effects of prenatal and postnatal tobacco smoke exposure on children. Archives of disease in childhood 88, 1086–1090 (2003).

9 Lange, S., Probst, C., Rehm, J. & Popova, S. National, regional, and global prevalence of smoking during pregnancy in the general population: a systematic review and meta-analysis. The Lancet. Global health 6, e769–e776, doi:10.1016/S2214-109X(18)30223-7 (2018).

10 Joehanes, R. et al. Epigenetic Signatures of Cigarette Smoking. Circulation. Cardiovascular genetics 9, 436–447, doi:10.1161/CIRCGENETICS.116.001506 (2016).

11 Joubert, B. R. et al. DNA Methylation in Newborns and Maternal Smoking in Pregnancy: Genome-wide Consortium Meta-analysis. American journal of human genetics 98, 680–696, doi:10.1016/j.ajhg.2016.02.019 (2016).

12 Richmond, R. C. et al. Prenatal exposure to maternal smoking and offspring DNA methylation across the lifecourse: findings from the Avon Longitudinal Study of Parents and Children (ALSPAC). Human molecular genetics 24, 2201–2217, doi:10.1093/hmg/ddu739 (2015).

13 Lee, K. W. et al. Prenatal exposure to maternal cigarette smoking and DNA methylation: epigenome-wide association in a discovery sample of adolescents and replication in an independent cohort at birth through 17 years of age. Environmental health perspectives 123, 193–199, doi:10.1289/ehp.1408614 (2015).

14 Richmond, R. C., Suderman, M., Langdon, R., Relton, C. L. & Davey Smith, G. DNA methylation as a marker for prenatal smoke exposure in adults. International journal of epidemiology, doi:10.1093/ije/dyy091 (2018).

15 Tehranifar, P. et al. Maternal cigarette smoking during pregnancy and offspring DNA methylation in midlife. Epigenetics 13, 129–134, doi:10.1080/15592294.2017.1325065 (2018).

16 Gaunt, T. R. et al. Systematic identification of genetic influences on methylation across the human life course. Genome Biol 17, 61, doi:10.1186/s13059-016-0926-z (2016).

17 Bonfiglio, F. et al. A GWAS meta-analysis from 5 population-based cohorts implicates ion channel genes in the pathogenesis of irritable bowel syndrome. Neurogastroenterology and motility : the official journal of the European Gastrointestinal Motility Society, e13358, doi:10.1111/nmo.13358 (2018).

18 Miettunen, J. et al. Data on schizotypy and affective scales are gender and education dependent‐‐study in the Northern Finland 1966 Birth Cohort. Psychiatry research 178, 408–413, doi:10.1016/j.psychres.2008.07.022 (2010).

19 Kupers, L. K. et al. DNA methylation mediates the effect of maternal smoking during pregnancy on birthweight of the offspring. International journal of epidemiology 44, 1224–1237, doi:10.1093/ije/dyv048 (2015).

20 Markunas, C. A. et al. Identification of DNA methylation changes in newborns related to maternal smoking during pregnancy. Environmental health perspectives 122, 1147–1153, doi:10.1289/ehp.1307892 (2014).

21 Lawlor, D. A. et al. Early life predictors of adolescent smoking: findings from the Mater-University study of pregnancy and its outcomes. Paediatric and perinatal epidemiology 19, 377–387, doi:10.1111/j.1365-3016.2005.00674.x (2005).

22 Taylor, A. E. et al. Maternal smoking during pregnancy and offspring smoking initiation: assessing the role of intrauterine exposure. Addiction 109, 1013–1021, doi:10.1111/add.12514 (2014).

23 Richiardi, L., Bellocco, R. & Zugna, D. Mediation analysis in epidemiology: methods, interpretation and bias. International journal of epidemiology 42, 1511–1519, doi:10.1093/ije/dyt127 (2013).

24 Hemani, G. et al. The MR-Base platform supports systematic causal inference across the human phenome. eLife 7, doi:10.7554/eLife.34408 (2018).

25 Ekblad, M., Gissler, M., Lehtonen, L. & Korkeila, J. Prenatal smoking exposure and the risk of psychiatric morbidity into young adulthood. Archives of general psychiatry 67, 841–849, doi:10.1001/archgenpsychiatry.2010.92 (2010).

26 Lahti, J. et al. Early-life origins of schizotypal traits in adulthood. The British journal of psychiatry : the journal of mental science 195, 132–137, doi:10.1192/bjp.bp.108.054387 (2009).

27 Niemela, S. et al. Prenatal Nicotine Exposure and Risk of Schizophrenia Among Offspring in a National Birth Cohort. The American journal of psychiatry 173, 799–806, doi:10.1176/appi.ajp.2016.15060800 (2016).

28 Talati, A. et al. Maternal smoking during pregnancy and bipolar disorder in offspring. The American journal of psychiatry 170, 1178–1185, doi:10.1176/appi.ajp.2013.12121500 (2013).

29 Larson, K. C., Lipko, M., Dabrowski, M. & Draper, M. P. Gng12 is a novel negative regulator of LPS-induced inflammation in the microglial cell line BV-2. Inflammation research : official journal of the European Histamine Research Society ... [et al.] 59, 15–22, doi:10.1007/s00011-009-0062-2 (2010).

30 Anderson, G. et al. Immuno-inflammatory, oxidative and nitrosative stress, and neuroprogressive pathways in the etiology, course and treatment of schizophrenia. Progress in neuropsychopharmacology & biological psychiatry 42, 1–4, doi:10.1016/j.pnpbp.2012.10.008 (2013).

31 Muller, N., Weidinger, E., Leitner, B. & Schwarz, M. J. The role of inflammation in schizophrenia. Frontiers in neuroscience 9, 372, doi:10.3389/fnins.2015.00372 (2015).

32 Khandaker, G. M., Zimbron, J., Dalman, C., Lewis, G. & Jones, P. B. Childhood infection and adult schizophrenia: a meta-analysis of population-based studies. Schizophrenia research 139, 161–168, doi:10.1016/j.schres.2012.05.023 (2012).

33 Aberg, K. A. et al. Testing two models describing how methylome-wide studies in blood are informative for psychiatric conditions. Epigenomics 5, 367–377, doi:10.2217/epi.13.36 (2013).

34 Rantakallio, P. The longitudinal study of the northern Finland birth cohort of 1966. Paediatr Perinat Epidemiol 2, 59–88 (1988).

35 Rantakallio, P. a. Acta Paediatr Scand 193, Suppl 193:191+ (1969).

36 Sovio, U. et al. Genetic determinants of height growth assessed longitudinally from infancy to adulthood in the northern Finland birth cohort 1966. PLoS Genet 5, e1000409, doi:10.1371/journal.pgen.1000409 (2009).

37 Jarvelin, M. R., Hartikainen-Sorri, A. L. & Rantakallio, P. Labour induction policy in hospitals of different levels of specialisation. British journal of obstetrics and gynaecology 100, 310–315 (1993).

38 Arshad, S. H. et al. Cohort Profile: The Isle Of Wight Whole Population Birth Cohort (IOWBC). International journal of epidemiology, doi:10.1093/ije/dyy023 (2018).

39 Boyd, A. et al. Cohort Profile: the ‘children of the 90s’‐‐the index offspring of the Avon Longitudinal Study of Parents and Children. International journal of epidemiology 42, 111–127, doi:10.1093/ije/dys064 (2013).

40 Fraser, A. et al. Cohort Profile: the Avon Longitudinal Study of Parents and Children: ALSPAC mothers cohort. International journal of epidemiology 42, 97–110, doi:10.1093/ije/dys066 (2013).

41 Relton, C. L. et al. Data Resource Profile: Accessible Resource for Integrated Epigenomic Studies (ARIES). Int J Epidemiol 44, 1181–1190, doi:10.1093/ije/dyv072 (2015).

42 Lehne, B. et al. A coherent approach for analysis of the Illumina HumanMethylation450 BeadChip improves data quality and performance in epigenome-wide association studies. Genome Biol 16, 37, doi:10.1186/s13059-015-0600-x (2015).

43 Houseman, E. A. et al. DNA methylation arrays as surrogate measures of cell mixture distribution. BMC Bioinformatics 13, 86, doi:10.1186/1471-2105-13-86 (2012).

44 Touleimat, N. & Tost, J. Complete pipeline for Infinium((R)) Human Methylation 450K BeadChip data processing using subset quantile normalization for accurate DNA methylation estimation. Epigenomics 4, 325–341, doi:10.2217/epi.12.21 (2012).

45 R: A language and environment for statistical computing (R Foundation for Statistical Computing, VIenna, Austria, 2017).

46 Taylor, A. E., Davey Smith, G., Bares, C. B., Edwards, A. C. & Munafo, M. R. Partner smoking and maternal cotinine during pregnancy: implications for negative control methods. Drug and alcohol dependence 139, 159–163, doi:10.1016/j.drugalcdep.2014.03.012 (2014).

47 Pinheiro, J. C. & Bates, D. M. Mixed-effects models in S and S-PLUS Springer. New York (2000).

48 Davey Smith, G. & Hemani, G. Mendelian randomization: genetic anchors for causal inference in epidemiological studies. Human molecular genetics 23, R89–98, doi:10.1093/hmg/ddu328 (2014).

49 Dusting Tingley, T. Y., Kentaro Hirose, Luke Keele, Kosuke Imai. mediation: R Package for Causal Mediation Analysis. . Journal of Statistical Software 59, 1–38 (2014).

50 Kosuke Imai, L. K., Tappei Yamamoto. Identification, Inference and Sensitivity Analysis for Causal Mediation Effects. Statistical Science 25, 51–71 (2010).

